# Investigating the impacts of sidechains on de-novo protein design

**DOI:** 10.1101/2025.08.08.669410

**Authors:** Cooper Svajda, Joshua Yuan

## Abstract

**De novo** protein design aims to create novel protein structures and sequences, often to enable specific functions. Most current generative models operate on simplified backbone-only representations. However, in vivo and in vitro protein folding and function are largely mediated by amino acid sidechains. Given this biochemical relevance, we ask: what happens when sidechain and sequence information are introduced into a protein generative model? To address this, we developed **SiBaSe**, a novel generative model that simultaneously co-designs sidechains, backbones, and sequence. SiBaSe achieves design performance approaching that of state-of-the-art backbone-only models. However, analysis of design patterns reveals that, despite access to sidechain data, the model behaves similarly to a backbone-based model. This appears to arise from uncertainties in the simultaneous modeling of sequence and sidechains that are inherent to flow-based architectures and offers new insight into architectural limitations and opportunities for improving generative protein design.

## 1 Introduction

Protein design is an emerging field with the potential to transform nearly every area of biology. By designing proteins *de novo* for specific functions, it becomes possible to bypass the constraints of natural evolution and create novel, non-natural phenotypes. Recent advances in this field have been driven by machine learning-based tools that automate and accelerate the design process.

The standard pipeline (Yeh et al. 2023, Lauko et al. 2024) begins with a **structure generation tool**, which produces a protein backbone lacking sequence or sidechain information (e.g., Watson et al. 2023, Yim et al. 2023a, Lin et al. 2023). These backbone structures can be guided toward specific functions through the inclusion of structural or functional motifs. Next, a **sequence design model**, most commonly ProteinMPNN (Dauparas et al. 2022), is used to infer an amino acid sequence intended to fold into the target backbone. Finally, the designed sequence is evaluated using a **structure prediction model**, such as AlphaFold2 or ESMFold, to assess whether the predicted fold matches the original design (Jumper et al. 2021, Lin et al. 2023). The agreement between the designed and predicted structures, often referred to as **designability**, is used to determine the in-silico success of the design.

This design pipeline has enabled the creation of novel and functional proteins that would have been unattainable only a few years ago, including *de novo* luciferases, serine hydrolases, and high-affinity binders with micro-to nanomolar affinities (Yeh et al. 2023, Lauko et al. 2023, Krishna et al. 2024). Remarkably, these successes have been achieved using models that, at a fundamental level, do not directly manipulate one of the most critical determinants of protein structure: the amino acid sidechains. According to Anfinsen’s Dogma, a protein’s sequence alone determines its folded structure, and the distinguishing chemical features of a sequence arise from the unique compositions and order of its sidechains (Anfinsen, 1973). Decades of biochemical research supports this, showing that folding is driven by sidechain interactions; whether through modulating the backbone’s electronic structure immediately after translation (Spassov et al. 2007) or later via sidechain packing, burial, and non-covalent interactions (Dill 1990, Dill et al. 2008, Farber & Mittermaier 2008). This biochemical importance stands in contrast to the simplifications of backbone-only models, prompting the central question of this study: **what happens when sidechain information is introduced into a protein generative model?**

To investigate this question, we developed a novel protein generative model called **SiBaSe** (Sidechains, Backbone, Sequence). Incorporating sidechains into the design process necessitated the inclusion of sequence information, as an amino acid’s identity is inherently defined by its sidechain. To isolate the effect of sidechain inclusion, we designed SiBaSe to be architecturally similar to a recent family of backbone models: FrameDiff, FrameFlow, and MultiFlow (Yim et al. 2023a, Yim et al. 2023b, Campbell et al. 2024), henceforth the FrameFamily. By maintaining architectural similarity with these models, we aimed to control for spurious variables and directly assess the impact of incorporating sidechain-level detail.

A key challenge in the co-generation of sidechains and backbones is the inherent sequence dependence of sidechains: their geometry and identity are tightly coupled. In *de novo* protein generation, however, the sequence is initially unknown, requiring a sidechain representation that is independent of residue identity. SiBaSe addresses this by representing sidechains using the same local-frame based system employed for the backbone. This choice enables the model to manipulate sidechain geometries without requiring prior knowledge of the sequence. Additionally, it facilitates seamless architectural integration, minimizes memory usage, and supports the use of the rotationally invariant Invariant Point Attention (IPA) module (Jumper et al. 2021).

To evaluate the impact of sidechain inclusion on generative performance, we compared SiBaSe to other models in the FrameFamily across three key metrics: **designability, diversity, and novelty**. To further probe the relative contribution of each data type (backbone, sidechain, and sequence) we conducted a **partial conditioning** experiment, selectively providing subsets of structural information during generation. Finally, we performed a qualitative assessment of model behavior, including **sequence confidence, sidechain placement**, and overall design patterns, to better understand how sidechain information influenced the generative process.

## 2 Results

### 2.1 Unconditional Generation

The first test of SiBaSe was the unconstrained design of 450 proteins ranging from 75 to 225 amino acids in length. Each of these generated proteins possessed a SiBaSe generated backbone structure, sequence and sidechain placements. The designed sequence of each protein was then provided to ESMFold (Lin et al. 2023), whose predictions were compared to the original designs to calculate designability (backbone-RMSD). Each of the outputs with designability <3 Å were then compared to their nearest structural neighbor in the PDB, using the program FoldSeek, which returned a pdbTM value for each design (VanKempen et al. 2023). SiBaSe was capable of designing protein structures and sequences which achieved designability backbone-RMSD values as low as 2.42 Å (Fig 1). Despite this, out of the 450 designs only 7 had designability scores of less than 3 Å (1.6%) This designability success rate is significantly lower than peer backbone-only models. It is important to note, the likelihood of successfully designing sequences and structures which fold at under 3 Å of accuracy by random chance is infinitesimal. Any level of success at this task indicates the model had learnt a true mapping in protein sequence-structure space.

**Figure 1:**
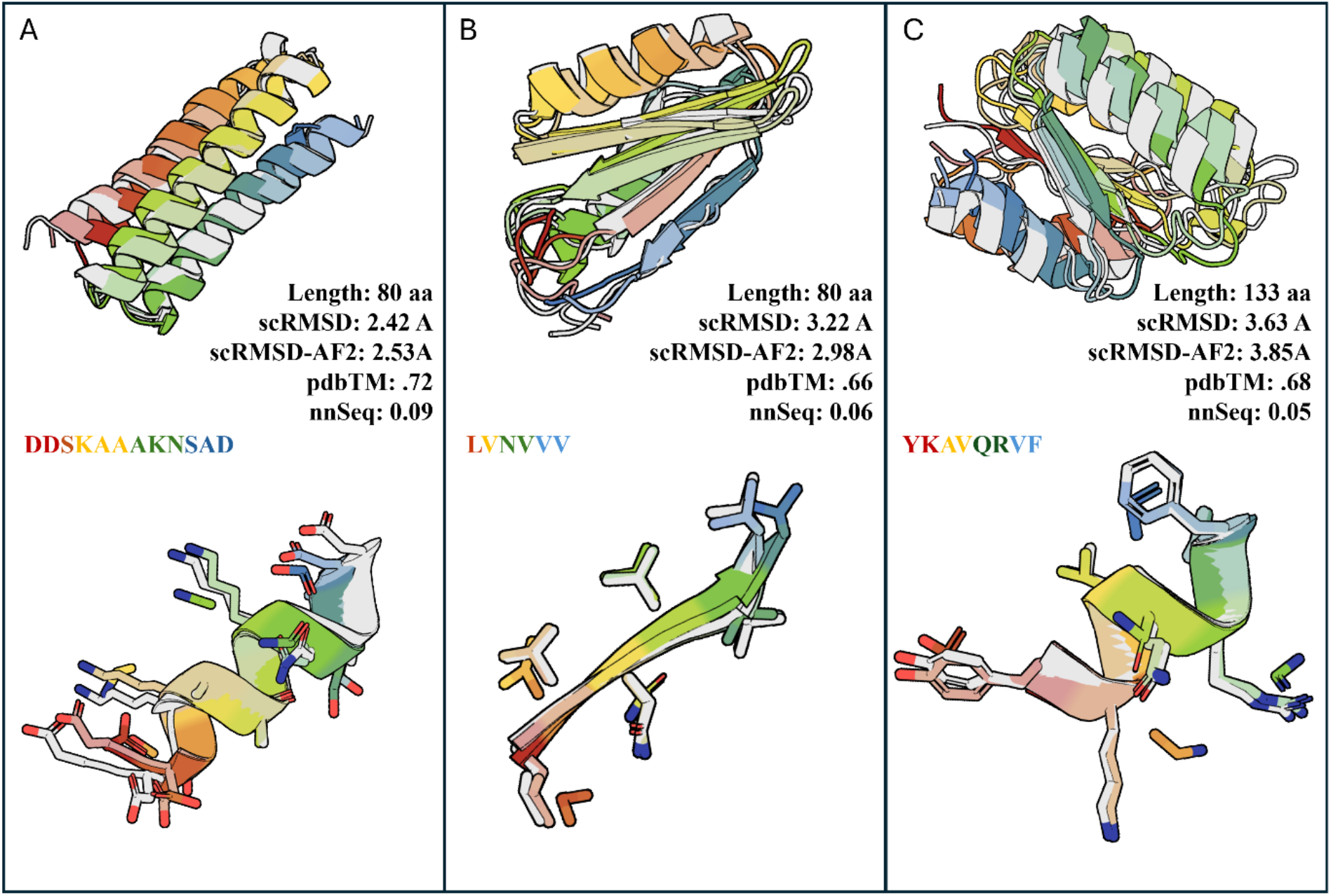
Unconditional Co-Generation Sampling of SiBaSe: Above are three proteins whose structure, sequence and sidechains were generated using SiBaSe. The saturated color is the SiBaSe output while the faded-colored and grey structures are the superimposed ESMFold and AlphaFold2 predictions respectively. The alignments are shown as **scRMSD** and **scRMSD-AF2** respectively. **pdbTM and nnSeq** record the TM score and sequence identity to the nearest neighbor in the PDB as determined by FoldSeek. The insets below show zoomed-in fragments with sidechain designs/predictions displayed in same color scheme as above. The sequence and corresponding fragment are color coded for amino acid identification.

In order to assess the quality of the protein structures themselves, and not the quality of sequence-structure pairs, the same 450 designs were each given 3 new sequences designed by ProteinMPNN (Dauparas et al. 2022). These sequences were then subjected to the same ESMFold reconstructions for designability assessment as before (Fig 2). Additionally, all designs with designability under 3 Å (the ‘success subset’) had a nearest-neighbor found in the PDB using FoldSeek with which to calculate pdbTM. The ‘success subset’ was then clustered via TM score following established conventions (Herbert 2008), with the number of clusters divided by total proteins in the subset to arrive at a diversity score.

**Figure 2:**
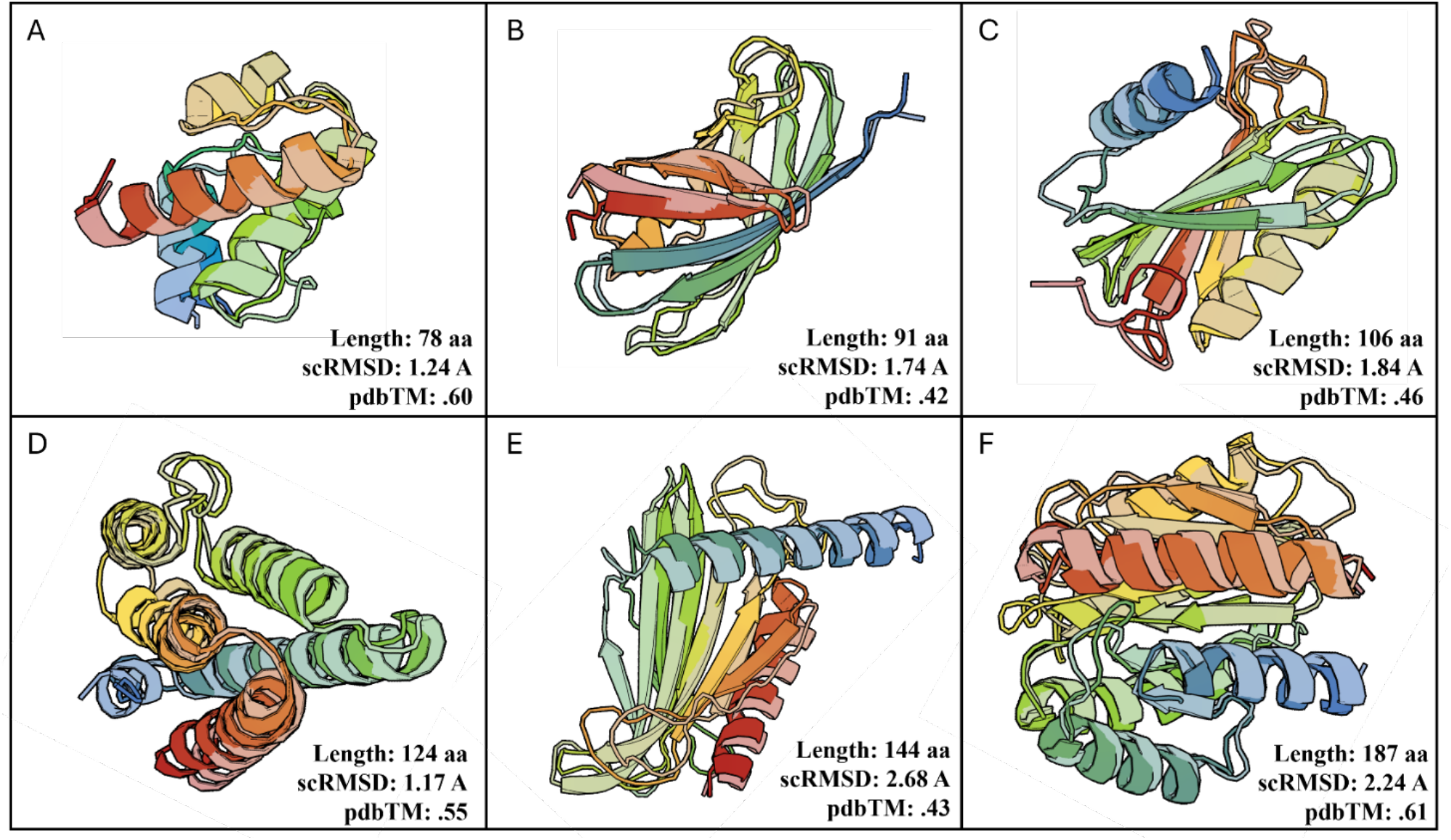
MPNN Redesigns of SiBaSe Designed Backbones: Above are 6 designs from SiBaSe whose sequences were redesigned using ProteinMPNN. The sequences were folded using ESMFold and structures aligned and visualized as in Figure 1. The scRMSD and pdbTM displayed derived from same approach described in Figure 1.

SiBaSe demonstrated generative behavior broadly consistent with other models in the FrameFamily across multiple key metrics (Table 1). Its generated structures were comparably The average diversity of its generated structures was comparable to that of Frame Diff and exceeded the diversity observed in FrameFlow. In the subset of successful designs, SiBaSe achieved slightly improved novelty relative to peer models. When evaluating designability, co-design success was observed in 1.6% of cases, a lower rate than backbone-only models, including the co-generative MultiFlow. However, when sidechains were excluded and structures were re-sequenced using ProteinMPNN, success rates improved substantially. At a 3 Å cutoff, designability reached 30%, and at the more stringent 2.5 Å cutoff used in prior FrameFamily publications, SiBaSe achieved a 24% success rate. Together, these results suggest that while co-design introduces challenges for structural fidelity, SiBaSe shares key generative characteristics with established backbone-focused models.

**Table 1:** Comparison of SiBaSe to FrameFamily: SDE – stochastic sampling strategy, ODE – deterministic/flow sampling strategy, CoDesign – simultaneous design of structure and sequence, * - proportion < 3 Å vs <2.5 Å for other designability values, **-value of 1 not reported due to overinflation from small number of total proteins, *** - no published values

| **Model** | **Steps** | **Method** | **Designability ↑** | **Diversity ↑** | **Novelty ↓** |
| --- | --- | --- | --- | --- | --- |
| FrameDiff | 500 | SDE | 0.42 | 0.36 | 0.66 |
| FrameDiff | 100 | SDE | 0.39 | 0.32 | 0.66 |
| FrameFlow | 500 | ODE | 0.81 | 0.22 | 0.69 |
| FrameFlow | 100 | ODE | 0.77 | 0.28 | 0.67 |
| MultiFlow | 500 | ODE | 0.89 | -*** | 0.62 |
| MultiFlow | 500 | ODE - codesign | 0.42 | -*** | 0.62 |
| SiBaSe | 250 | SDE | 0.30 * | 0.31 | 0.60 |
| SiBaSe | 250 | SDE - codesign | 0.016 * | - ** | 0.69 |

In nearly all cases, sequences redesigned with ProteinMPNN achieved higher designability than the original SiBaSe-generated sequences (Fig. 3A). While SiBaSe was capable of producing successful designs, a gradual decline in designability was observed with increasing protein length (Fig. 3B). Although novelty showed a modest negative correlation with designability, the model was still able to generate proteins that were both highly novel and structurally accurate (Fig. 3C). In terms of secondary structure composition, SiBaSe exhibited no clear bias, generating proteins with diverse secondary structural composition (Fig. 3D).

**Figure 3:**
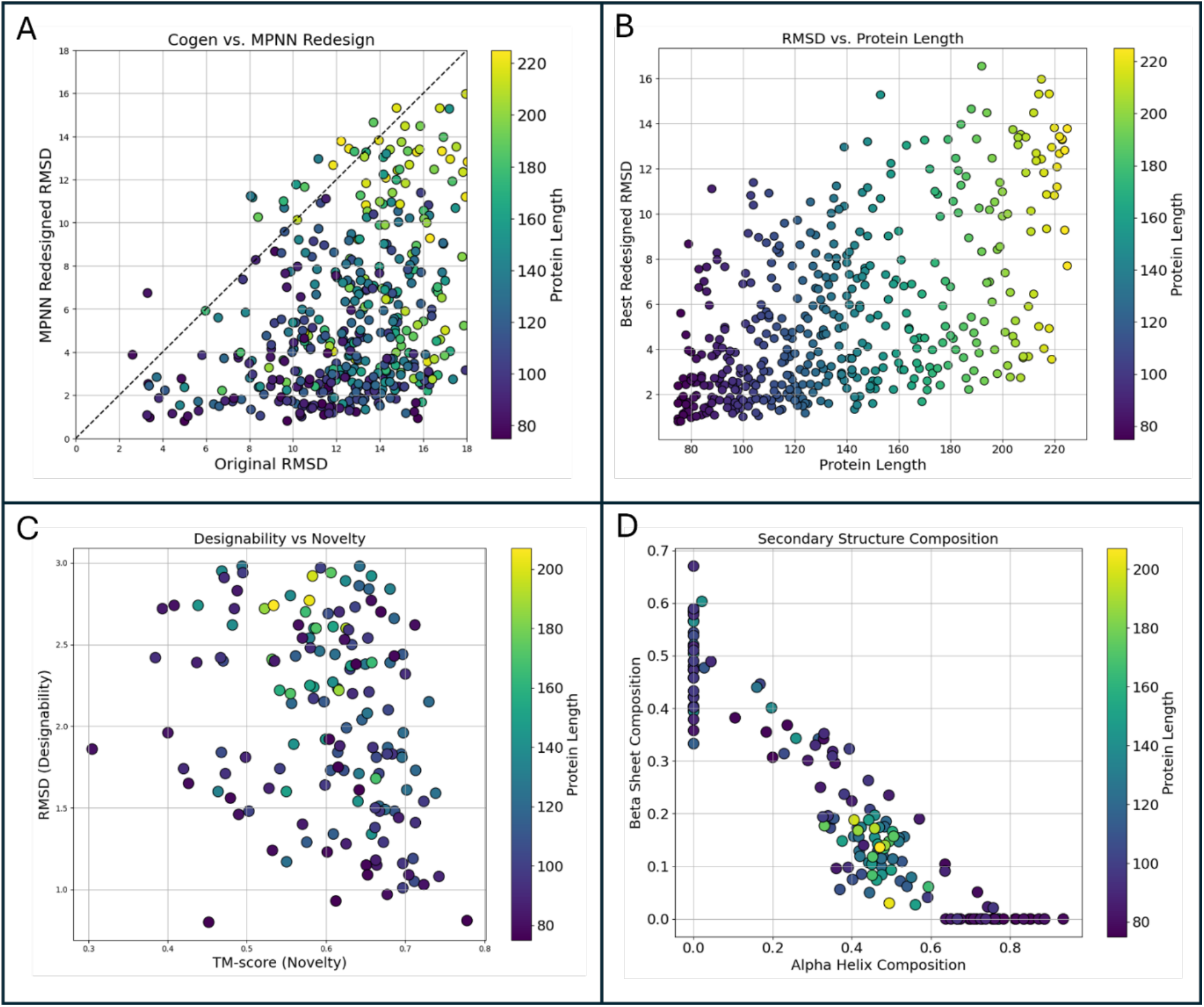
Analysis of SiBaSe Designs: Graphical representation of SiBaSe designs illustrating designability versus sequence origin, length and novelty. An additional panel illustrates secondary structure diversity of designs. Panel A plots designability RMSD for SiBaSe designed sequence (x-axis) versus MPNN redesign (y-axis). Panel B plots designability RMSD (bester performer, y-axis) against protein length (x-axis). Panel C plots designability RMSD (y-axis) against novelty (TM, x-axis). Panel D plots proportion of secondary structure; y-axis is beta-sheet proportion, x-axis is alpha-helix proportion.

### 2.2 Conditional Generation

In an attempt to investigate the relative impact or importance of backbone and sidechain geometric information during design, an experiment was conducted using conditioning. Briefly, conditioning involves providing the model with a fragment(s) of protein structure and/or sequence called the ‘motif’ and tasking it with generating a protein containing it, the ‘scaffold’ (Fig 4). The replacement method of conditioning, while facing key limitations, was more than adequate for this experiment and required no specialized training for the conditioning task (Trippe et al. 2022).

**Figure 4:**
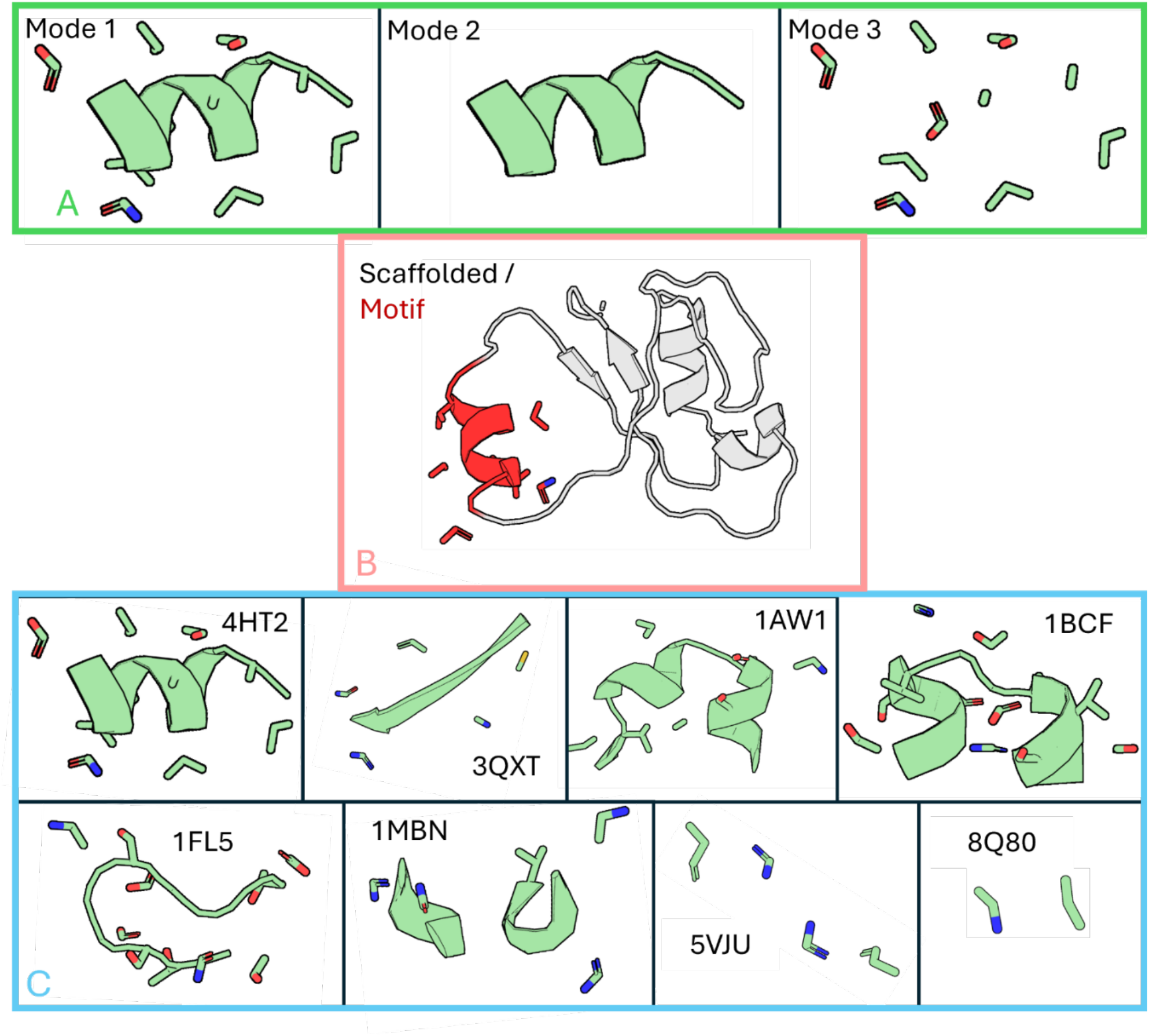
Conditioning Modes and Motifs: Above is a graphical representation of the 3 conditioning modes used in this work, the concept of conditioning, and the 8 motifs used in this work. Panel A shows three conditioning modes: 1 – backbone/sidechain/sequence, 2 – backbone/sequence, 3 – sidechain/sequence. Panel B illustrates the process of conditioning, motif in red scaffold in grey. Panel C shows the 8 motifs used, the PDB ID of origin for each motif is shown next to the motifs.

To evaluate how different types of structural information influence design, we tested SiBaSe under three distinct conditioning modes. In **Mode 1**, the conditioning motif included backbone, sidechains, and sequence. **Mode 2** used backbone and sequence, while **Mode 3** provided sidechains and sequence without the backbone (Fig 4A). Motifs were sampled from the Protein Data Bank (PDB) and selected to present increasing levels of difficulty: ranging from large, contiguous secondary structure elements to small, non-contiguous fragments. For each mode– motif combination, 30 designs were generated (Fig 4b). Each SiBaSe-designed sequence, along with three ProteinMPNN redesigns (with the motif sequence fixed), was submitted to ESMFold for structure prediction. Design–prediction pairs were evaluated using whole-protein designability scores as before, and additionally, **motif-only RMSD** was computed in an all-atom manner to assess local structural fidelity.

Designability scores, both for whole-protein structures and motif-level all-atom RMSD, declined with increasing motif complexity and across conditioning modes, with Modes 1 and 2 consistently outperforming Mode 3 (Fig. 5A). Because RMSD is sensitive to the size of the structure being evaluated, motif-level RMSDs were normalized to allow fair comparison across motifs of varying sizes. This normalization accentuated the observed trend, reinforcing the correlation between reduced motif size, or lack of backbones, and lower design accuracy (Fig. 5B). These results suggest that the model more effectively integrates backbone information than sidechain information during the generative process. Moreover, relatively small amounts of backbone input were sufficient to guide design outcomes, whereas equivalent sidechain information had a more limited effect (Fig. 5B).

**Figure 5:**
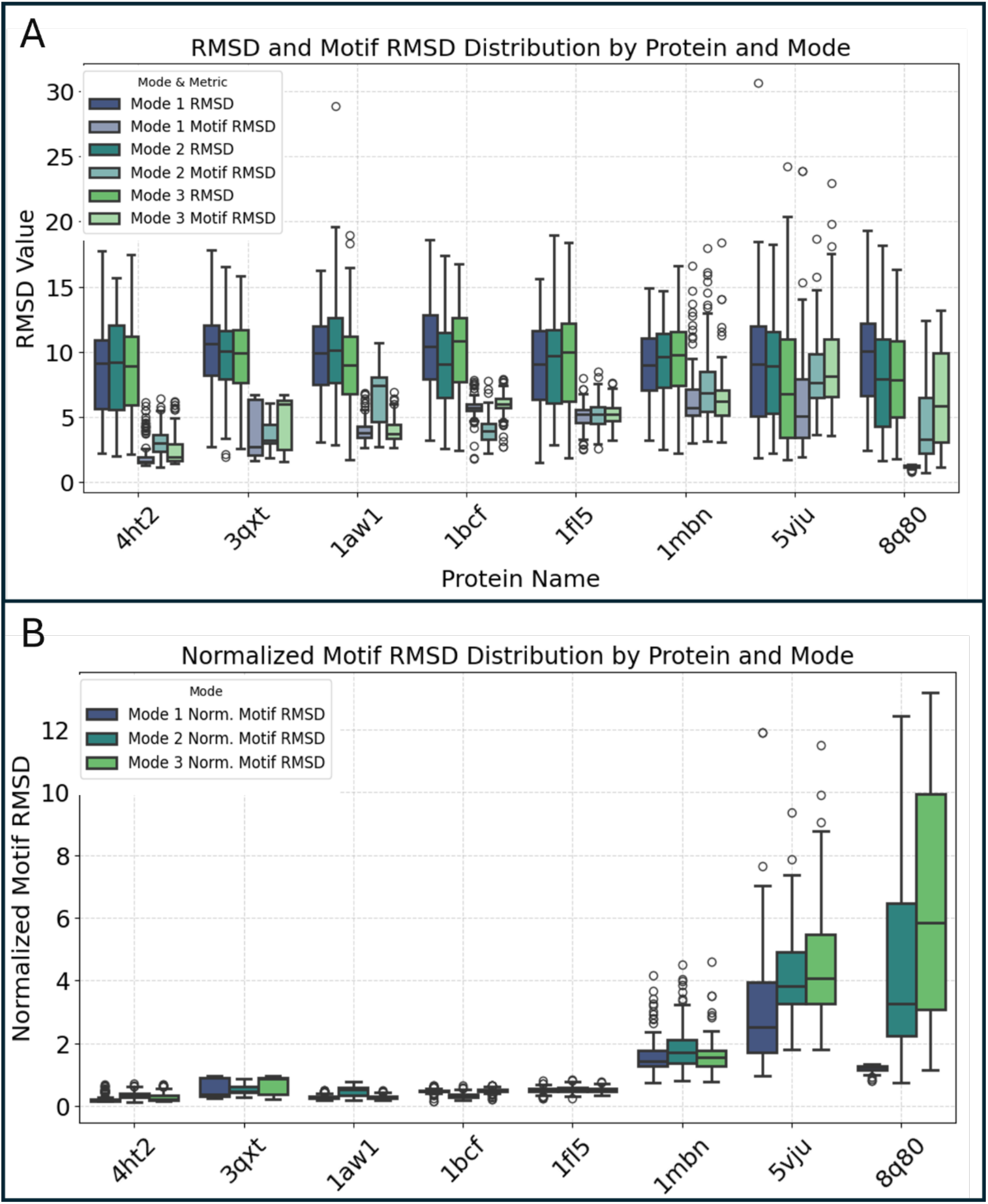
Conditional Sampling Evaluation: Above is a graphical representation, via boxplots, of the designability of motif-conditioned outputs of SiBaSe. Panel A shows the 8 motifs (x-axis) in order of increasing ‘difficulty’ from left to right. Each motif has 6 boxplots (left to right) corresponding to whole protein RMSD, modes 1, 2 and 3, then motif-all-atom RMSD, modes 1, 2, 3. Panel B shows the normalized motif-all-atom RMSD for the 8 motifs, 3 boxplots correspond (left to right) modes 1, 2, 3.

### 2.3 Sidechain Placement

Sidechain placement was analyzed by comparing 450 unconditioned SiBaSe designs to 450 randomly sampled proteins from the training set. Amino acids were separated by type, and their backbone frames were aligned to assess sidechain translation. The central atom of each sidechain frame was plotted, and normalized Earth Mover’s Distance (nEMD) was computed to quantify similarity between real and designed point clouds (Fig. 6A). Placement accuracy varied by amino acid type: smaller, more common, and less flexible residues (e.g., Leucine, Asparagine, Valine, Glutamate) showed tighter cluster overlap and lower nEMD values, while larger, flexible, and rarer residues (e.g., Arginine, Methionine, Lysine) diverged more. A key observation is that the placement of designed sidechains was slightly closer to the backbone as compared to the real protein distributions for most of the amino acid types (exceptions being Valine and Leucine). Serine and Isoleucine were notable exceptions to this pattern with poor performance despite their size and frequency in the training data.

**Figure 6:**
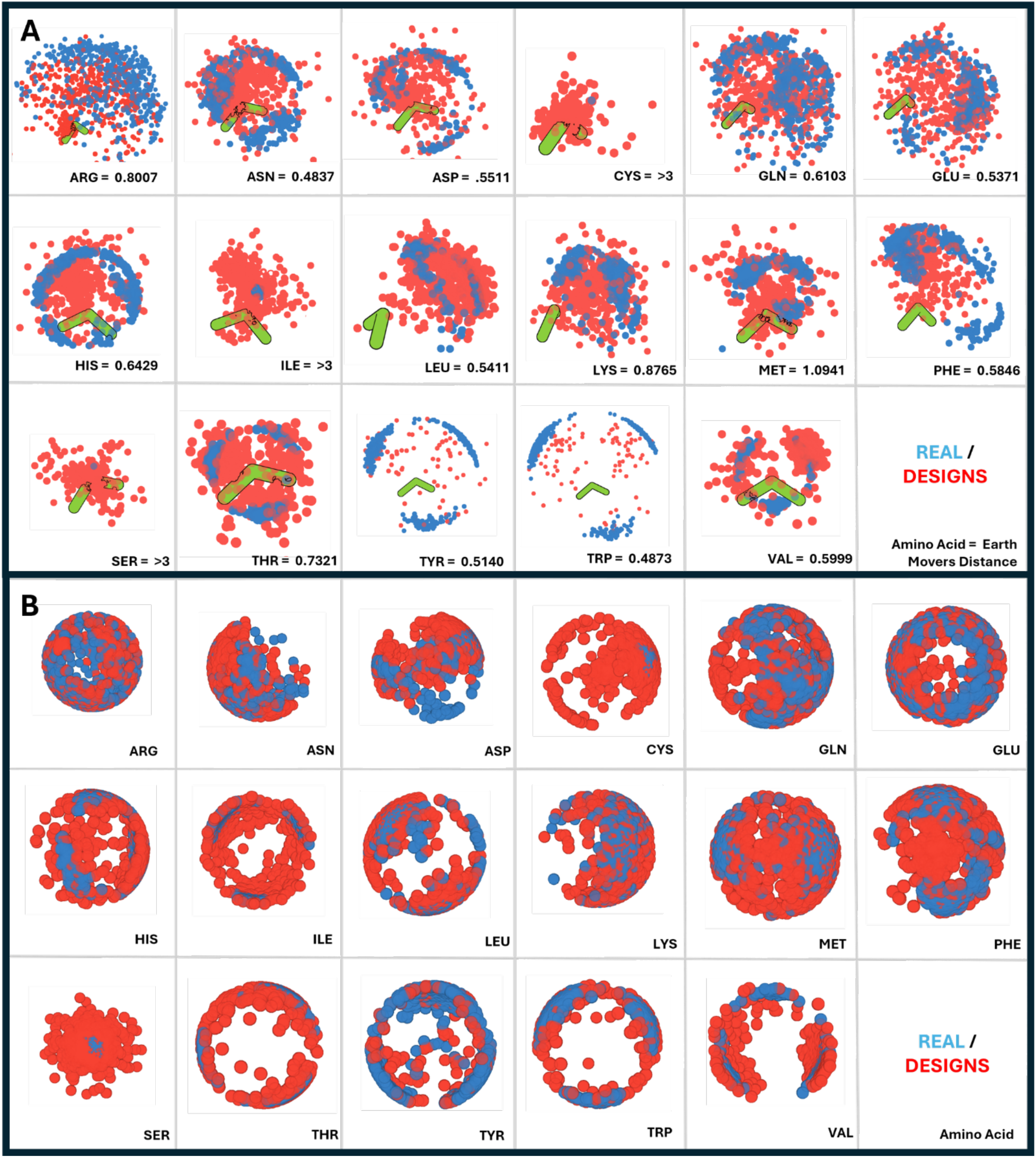
Sidechain Placement Visualization: Above is an illustration of the sidechain frame placements in SiBaSe designs versus real proteins. Panel A demonstrates the sidechain placement (translation) of real (blue) and designed (red) amino acids relative to the amino acid backbone (green). The normalized earth movers distance (nEMD) value is adjacent the amino acid name. Panel B shows the orientations of amino acids sidechains in a unit-sphere-like representation. Real sidechain frame terminal atoms are shown in blue, designs shown in red.

To assess orientation, the same central atoms were centered at the origin and the terminal frame atoms were projected onto a unit sphere (Fig. 6B). Orientation accuracy similarly depended on residue type, with narrow, well-overlapping distributions for Tyrosine, Tryptophan, Valine, Isoleucine, and Threonine, and broader divergence for Arginine, Methionine, and Lysine. Interestingly, Serine and Isoleucine again showed strong orientation overlap despite translational inaccuracy.

### 2.4 Sequence Generation Dynamics

Following the continuous-time Markov chain framework described in MultiFlow (Campbell et al. 2024), SiBaSe outputs a rate matrix that reflects the model’s confidence across all 20 amino acid types at each position during generation. This enables direct visualization of sequence dynamics throughout the design process (Fig. 7). Initially, with the sequence tensor fully masked, the model exhibits low confidence across all positions, as indicated by low color saturation. Around timestep *t* = 0.33, confidence begins to increase and the model starts converging on a prospective sequence. However, even after this point, many positions continue to undergo rapid turnover in predicted amino acid identity. The most confident residue at a given position often shifts significantly in both biochemical properties and size. One illustrative example (highlighted by a red arrow in Fig 7) shows a single position transitioning from alanine to phenylalanine, cysteine, glutamate, lysine, and finally leucine over the course of a single trajectory.

**Figure 7:**
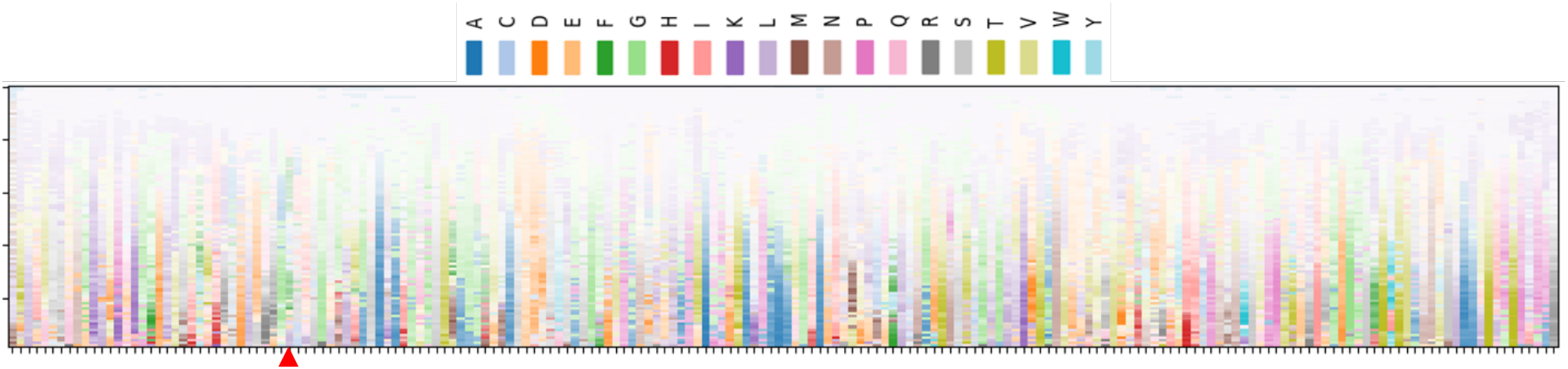
Amino Acid Confidence Matrix: The above matrix visualizes the amino acid sequence, and confidence, of a SiBaSe generated protein during the design process. X-axis corresponds to the amino acid sequence of designed protein in N-to-C order (left to right). Y-axis corresponds to generative timestep, top of graph *t* = 0, bottom of graph *t* = 1. Color corresponds to amino acid type. Saturation indicates confidence, lower saturation indicating lower confidence.

### 2.4 Intra-Generation Structural Dynamics

As a final assessment of model behavior during the generative process, a single design trajectory was sampled, and the model’s outputs were visualized at each timestep (Fig. 8). Protein backbone frames were rendered in purple and sidechain frames in green. In the final structure, sidechain frames were well-distributed, filling the space around and between backbone elements. Earlier in the trajectory (*t* < 0.75), however, sidechain frames remained closely clustered around their respective backbone frames. The backbone appeared to extend and adopt a coarse fold first, with sidechain frames remaining tightly coupled to their backbones. This visualization highlights a temporal separation in structure formation, with backbone geometry evolving earlier in the trajectory and sidechain placement following.

**Figure 8:**
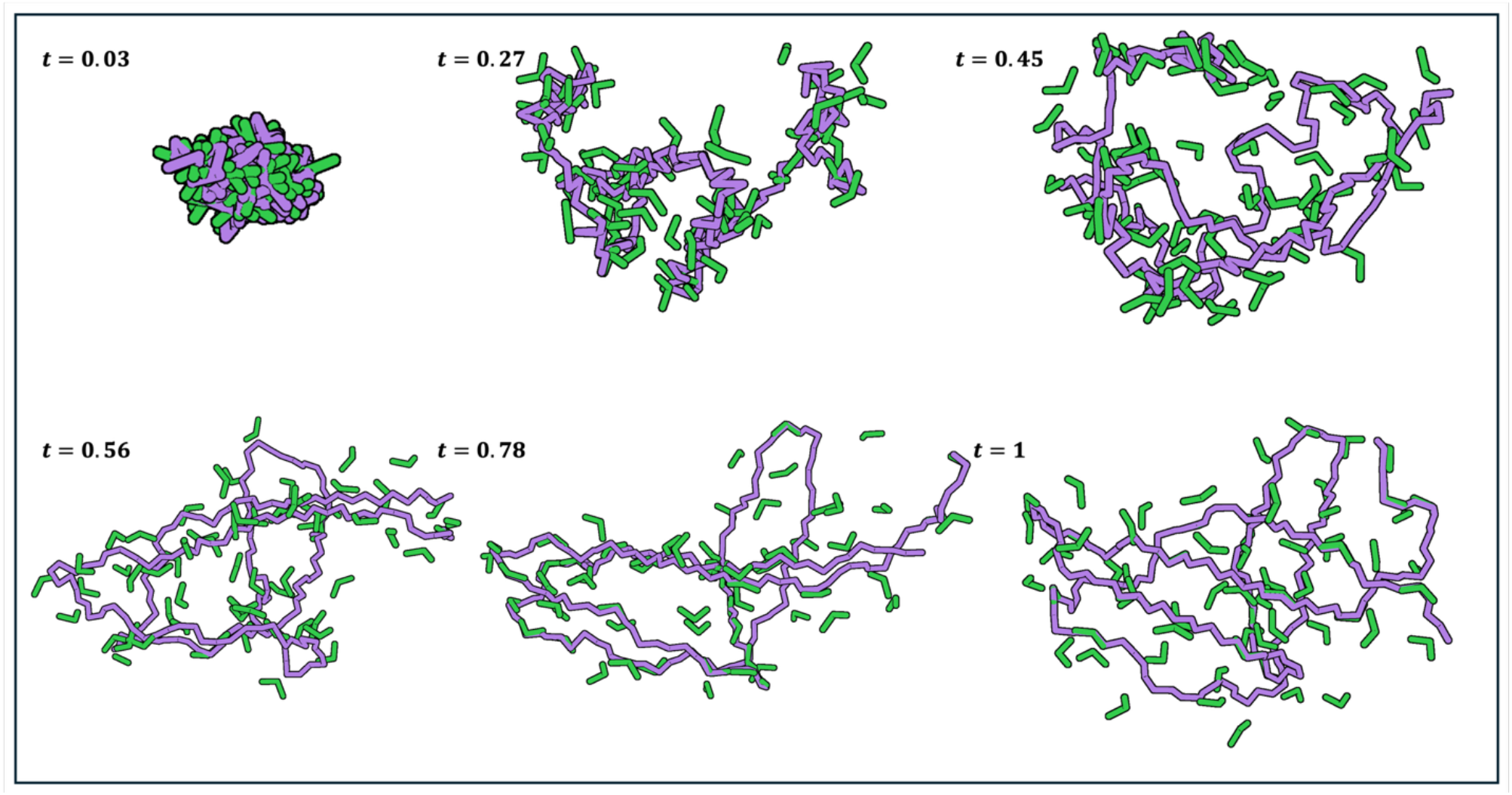
Visualization of Protein Generative Trajectory within SiBaSe: Above are 6 snapshots of the SiBaSe protein ‘prediction’, or output, for 6 timesteps during a single design trajectory. Timestep value of snapshot to the top left of each image. Backbone frames are colored purple and sidechain frames colored green.

## 3 Discussion

Sidechains play an indispensable role in protein folding and function (Dill 1990, Dill et al. 2008, Spassov et al. 2007, Farber & Mittermaier 2008), so their inclusion in a generative model was expected to produce significant deviations in behavior from backbone-only structural models. However, the most immediate and striking result was that SiBaSe, despite incorporating sidechain information, exhibited only modest differences from the backbone-focused FrameFamily models in terms of designability, novelty, and diversity. The co-generation success rate of 1.6% (at a <3 Å threshold) was notably lower than that of MultiFlow, the only other model capable of joint sequence and structure generation. Nonetheless, even modest success in the co-design task demonstrates that SiBaSe was able to learn a non-trivial mapping between structure and sequence spaces. When structure alone was evaluated, by remaking sequence with ProteinMPNN, SiBaSe achieved a substantially higher success rate of 30%, revealing a stark contrast between structure-only and co-generation performance. This divergence in structure-only vs cogeneration success pointed toward a deeper phenomenon that was further explored through targeted analyses.

In order to assess the relative contributions of the various kinds of information, conditioning was used as a method for controllable data-injection. During the design process, the positions/sequence of the motif inform the surrounding designed protein, and as such the models ability to utilize this information serves as a practical proxy for the importance and interpretability of the injected data. As expected, designability was highest for larger and structurally simpler motifs. However, as motif size decreased, the performance of sidechain-only (Mode 3) outputs declined more rapidly than that of backbone-only (Mode 2) outputs. Visual inspection of selected designs (Supp. Fig. 3) revealed that small sidechain-only motifs were often disregarded entirely, with final sidechain frames appearing up to 10 Å away from their corresponding backbone positions. This suggests that the model had more difficulty extracting actionable structural cues from sidechains alone, particularly in sparse or fragmentary contexts.

Generative models generate from distributions they have learned, so visualizing the placements of sidechains (tantamount to rotamer assignment) was expected to yield clues about the learning that took place in sidechain space. In terms of translational accuracy, the model was generally able to assign plausible sidechain positions relative to the backbone, with performance correlated to amino acid size, frequency, and flexibility. However, the predicted sidechains tended to remain closer to the backbone than those in real proteins, suggesting a conservative placement strategy. Notably, Serine and Isoleucine exhibited unexpectedly poor translational accuracy despite their high frequency and simple rotameric profiles. In contrast, orientation accuracy was more consistent across residue types. This pattern may reflect differences in learning complexity, as the space of orientations (the *SO*(3) manifold) is more constrained and potentially easier to model than the open-ended Cartesian space of translations. Overall, the model showed substantial divergence in sidechain placement accuracy between amino acid types and had an overall pattern of timid translational placement.

Sequence design in SiBaSe, following the continuous-time Markov chain approach of FrameFlow, proceeds via iterative updates of amino acid type probability distributions. A key advantage of this framework is that it enables inspection of the model’s internal decision-making dynamics during generation. Early in the design process, the model exhibited high uncertainty across the sequence, with low confidence in amino acid identity for the first third to half of the trajectory. Even after convergence began, substantial stochasticity remained, both in confidence levels and in the identity of the most probable amino acid at each position. In many cases, the most confident residue type at a given site shifted drastically between chemically and structurally distinct amino acids, suggesting ongoing competition between diverse sequence solutions.

These patterns come into focus with the last data point, the visualization of a generative trajectory. The models’ behavior was backbone-centric, in spite of the sidechain information present. For the majority of the trajectory, the backbone was assuming a nascent-then-final shape while the sidechains were ‘dragged’ along by their respective backbones. Rather than any apparent sharing of burden between backbones and sidechains with respect to structure resolution, the model appears to have learned to be a backbone-first model with sidechains serving a supporting role at best; and potentially a drag on the model’s design freedom at worst.

In answering why, we postulate that there is a ‘penalty of uncertainty’ which is a result of the interplay between sequence and sidechain geometry. Throughout the design trajectory, sequence predictions remained volatile, and sidechains were often positioned conservatively, closer to the backbone than observed in real proteins. While changes in sequence do not necessarily perturb backbone geometry, each shift in amino acid identity implies a distinct sidechain conformation, often requiring substantial spatial rearrangement. This geometric sensitivity introduces inertia into the design process: the model defers confident sidechain placement until the backbone stabilizes. As a result, sidechains are treated as secondary elements, trailing behind backbone formation rather than serving as active guides.

As a result, SiBaSe operates as a backbone-first model, with sidechains integrated late in the process and contributing minimally to early structural decisions. This highlights a key architectural limitation: without explicit mechanisms to resolve or mitigate the uncertainty from sidechain-sequence interdependence, the generative process defaults to the simpler, and more stable, backbone space.

## 4 Conclusions

SiBaSe was developed to explore the effect of incorporating biochemically rich sidechain information into generative protein design, and it was able to achieve near-peer performance in several metrics. Experimentation with the model revealed that despite having access to sidechain data, the model ultimately adopted a backbone-centric strategy. This suggests that the presence of sidechains, when represented in a singular format, introduces uncertainty that the model is reluctant to resolve. This behavior is likely generalizable to other computational approaches that treat sidechains deterministically during sequence generation. To meaningfully leverage the informational richness of sidechains, future models may need to adopt representations that accommodate multiple evolving sidechain configurations in parallel. Such flexibility could enable more effective exploration of the intertwined sequence–structure landscape and help overcome the architectural inertia imposed by sidechain uncertainty.

## 5 Materials and Methods

### 5.1 Flow Models

SiBaSe was designed to emulate the behavior of FrameFlow/MultiFlow, and as such it is of the Continuous Normalizing Flow (CNF) family of models (Yim et al. 2023a, Cambell et al. 2024). These models learn a reversible mapping between two distributions, the data distribution, *p*_1_, and some prior distribution (often a form of noise), *p*_0_, by integrating an ordinary differential equation (ODE) over a learned vector field,, *v*_θ_.

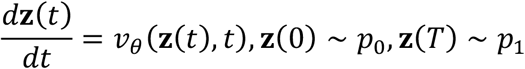

Flow matching is an approach to training in which, rather than solving the entirety of the differential equation, the model learns an approximation of the full vector field, *u*_*t*_ (*x* ∣ *x*_1_), in a supervised fashion (Lipman et al. 2022).

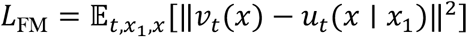

Progress along the vector field is quantified by a continuous variable, the timestep (*t*), in the range of [0, 1]. The noise priors exist at *t* = 0, and true data at *t* = 1. Intermediate levels of noise in the range [0, 1] are arrived at via a linear interpolation along the vector connecting the two points in either Euclidian space for coordinates, or the geodesic on the *SO*(3) manifold for orientations (Supp. Fig. 1). The selected prior distributions are a three-dimensional gaussian distribution, *N*(0,1) ∈ ℝ^3^, and the uniform distribution in *SO*(3), *U*(*SO*(3)), for coordinate and orientation data respectively. Sequence, as categorical data, will be described in section 5.3.

### 5.2 Frame Representation of Backbones and Sidechains

The backbones and sidechains of amino acids are represented as frames, mathematical objects with a coordinate and rotation matrix which describes a local reference frame. Frames are constructed, using the Gram-Schmidt orthogonalization process, from 3 points in space. Backbones are readily convertible in this format as they consist of a repeating pattern of N-C-Cα atoms with consistent internal bond geometry.

Representing sidechains as frames has several benefits, namely the sequence-agnosticism of the frame representation and ease of incorporating these additional frames with those of the backbones. In order to convert the sidechains into a frame representation it was necessary to create a library of atom types for each amino acid type which met 3 criteria:

‐ The atoms needed to possess the biochemically relevant atoms (ex. Charged atoms)
‐ The atoms needed to be part of a rigid group for un-orthogonalization
‐ The atoms needed to be distal to the backbone to capture full sidechain size

The atoms which were determined best to meet the listed criteria are shown in Table 2.

**Table 2:** Library of Amino Acid Sidechain Atoms for Frame Construction. The table above shows the library of atoms that would be used, for each amino acid type, to construct its sidechain frames. For amino acids with ‘NA’, the sidechain is either non-existent or exists as a rigid group with no rotamers. In these cases, the backbone frame was duplicated to serve as a placeholder.

| Amino Acid | Framing Atoms | Amino Acid | Framing Atoms |
| --- | --- | --- | --- |
| Alanine | NA/NA/NA | Leucine | CD1/CG/CD |
| Arginine | NH1/CZ/NH2 | Lysine | CD/CE/NZ |
| Asparagine | ND2/CG/OD1 | Methionine | CG/SD/CE |
| Aspartate | OD1/CG/OD2 | Phenylalanine | CE1/CZ/CE2 |
| Cysteine | CA/CB/SG | Proline | NA/NA/NA |
| Glutamine | NE2/CD/OE1 | Serine | CA/CB/OG |
| Glutamate | OE1/CD/OE2 | Threonine | OG1/CB/CG2 |
| Glycine | NA/NA/NA | Tryptophan | NE1/CE2/CZ2 |
| Histidine | ND1/CE1/NE2 | Tyrosine | OH/CZ/CE1 |
| Isoleucine | CB/CG1/CD1 | Valine | CG1/CB/CG2 |

### 5.3 Categorical Data

Categorical data is not readily usable in its native format within a flow model. As such, SiBaSe take the continuous-time Markov chain approach from MultiFlow (Campbell et al. 2024). In this approach, the sequence of the protein is represented as a one-hot encoded tensor for all amino acid types and masking tensor. The model predicts a rate matrix, or confidence matrix, of each amino acid type for each position. Further description below.

### 5.4 SiBaSe Architecture, Training and Sampling

SiBaSe’s training dataset consisted of 3902 proteins from the Protein Data Bank (PDB) that were monomeric, soluble proteins ranging from 75-225 amino acids in length, post-screened to remove proteins with missing loops/fragments. For each training step, SiBaSe would sample a protein from the training set, a timestep value *t*, and matching-size noise tensors from the *N*(0,1) ∈ ℝ^3^and *U*(*SO*(3)) distributions (for coordinate and rotations respectively). The timestep was scaled for rotations so that any *t* < 0.1 → *t* = 0. Finally, an independent timestep was sampled for the sequence.

The geometric data was interpolated to the selected noise levels following the mathematical approach in FramFlow (Yim et al. 2023b). The sequence timestep was then used to guide the random masking of sequence information such that the proportion of masked positions approximated (1 − *t*) as in MultiFlow. The final noised protein was passed to the model to generate a prediction for the original un-noised protein, 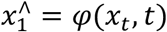. The model’s loss was tripartite, with the *SE*(3) loss of the FrameFamily (ℒ_*SE*3_), a cross-entropy loss on sequence (ℒ_*seq*_), and pairwise distance topology loss (ℒ_*topology*_), being scaled by coefficients and then combined to produce the final loss, ℒ_*total*_. The topology loss was a MSE loss on pairwise distances, with separate distance cutoffs for backbone-backbone (12 Å), backbone-sidechain (8 Å) and sidechain-sidechain (8 Å) components derived from the true structure distograms.

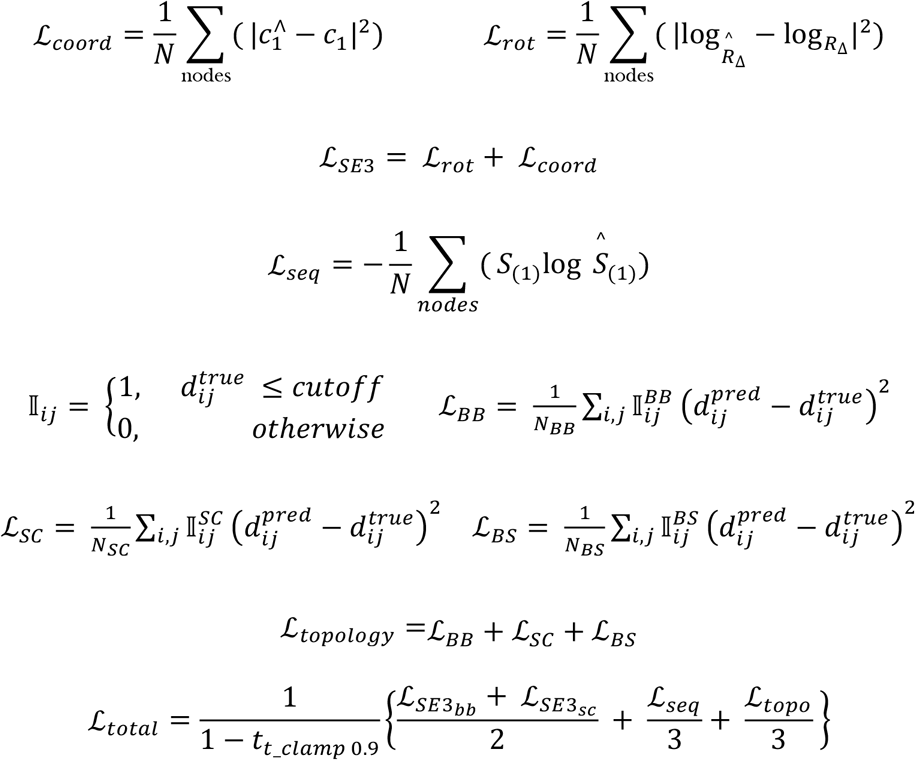

SiBaSe was trained for 28 days (250 epochs) on a single T4 GPU using the Adam Optimizer (Kingman and Ba, 2014), with a learning rate of 1e-4, β_1_=0.9 and β_2_=0.999. Each training set sample was drawn singly and then duplicated along the batch dimension, with independent timesteps and noise between batches, in a size-dependent manner to avoid memory waste on padding. After each of the final 80 epochs, the model generated 30 proteins for designability quantification with ESMFold. The checkpoints of the top 4 epochs (lowest designability score) were retained and the checkpoint from epoch 239 was selected as the vinal version (Supp. Fig. 4).

To sample SiBaSe, the model begins by drawing from the prior described noise distributions (*N*(0,1) ∈ ℝ^3^ and *U*(*SO*(3))) to generate starting tensors of the desired protein length. A corresponding fully-masked sequence tensor is generated. The model generates a prediction for the true protein, 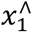, at each step. Initially, 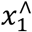 was simply interpolated back to the next timestep using the ODE approach of FrameFlow/MultiFlow. It was discovered that the model would settle on sub-optimal solutions (backbone breaks and clashes) earth in design and not be capable of escaping. To circumvent this, a stochastic approach was adopted. In the new approach, each time the model generates a prediction 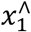, new noise tensors are sampled, and the prediction is noised using these new tensors to the next timestep input. In doing this, the model was capable of escaping early suboptimal solutions.

Following FrameDiff/FrameFlow/MultiFlow, SiBaSe is constructed from repeating layers of invariant point attention (IPA), MLP, Transformer-encoder and Geometric Update layers (Supp. Fig. 2). The main difference between SiBaSe and the models in the FrameFamily is the absence of edge representation. This is due to memory constraints that were created after the addition of sidechain frames, which effectively double the protein size. The model hyperparameters are as follows:

|  |  |  |  |
| --- | --- | --- | --- |
| Global Parameters: | node dim = 512 | L = 4 |  |
| IPA Parameters: | heads = 8 | query points = 8 | value points = 12 |
| Transformer Parameters: | heads = 8 | layers = 4 |  |

SiBaSe has a total of 33.8 million trainable parameters.

